# Monsters with a shortened vertebral column: A population phenomenon in radiating fish *Labeobarbus* (Cyprinidae)

**DOI:** 10.1101/2020.09.11.292763

**Authors:** Alexander S. Golubtsov, Nikolai B. Korostelev, Boris A. Levin

## Abstract

The phenomenon of a massive vertebral deformity was recorded in the radiating *Labeobarbus* assemblage from the middle reaches of the Genale River (south-eastern Ethiopia, East Africa). Within this sympatric assemblage, five trophic morphs – generalized, lipped, piscivorous and two scraping feeders – were reported between 1993 and 2019. In 2009, a new morph with prevalence of ∼10% was discovered. The new morph, termed ‘short’, had an abnormally shortened vertebral column and a significantly heightened body. This type of deformity is common in farmed Atlantic salmon and other artificially reared fish, but is rare in nature. In the Genale *Labeobarbus* assemblage, the deformity was present exclusively within the generalized and lipped morphs. The short morph had between seven and 36 deformed (compressed and/or fused) vertebrae. Their body height was positively correlated with number of deformed vertebrae. In another collection in 2019, the short morph was still present at a frequency of 11%. Various environmental and genetic factors could contribute to the development of this deformity in the Genale *Labeobarbus*, but based on the available data, it is impossible to confidently identify the key factor(s). Whether the result of genetics, the environment, or both, this high-bodied phenotype is assumed to be an anti-predator adaptation, as there is evidence of its selective advantage in the generalized morph. The Genale “monstrosity” is the first reported case of a massive deformity of the vertebral column in a natural population of African fishes.

“We have also what are called monstrosities; but they graduate into varieties. By a monstrosity I presume is meant some considerable deviation of structure in one part, either injurious to or not useful to the species, and not generally propagated. If it could be shown that monstrosities were even propagated for a succession of generations in a state of nature, modifications might be effected (with the aid of natural selection) more abruptly than I am inclined to believe they are.” Darwin (1860, pp. 46, 426).

## INTRODUCTION

The emergence and establishment of morphological novelties in a population is central to morphological evolution, although this process remains poorly understood. As initially highlighted by Darwin [1], there is no clear discrimination between morphological abnormalities (monstrosities) and regular variation. Morphological abnormalities that are caused by genetic factors – and hence may be shaped by natural selection – are particularly interesting to evolutionary biologists as potential novelties.

Ray-finned fishes form one of the major vertebrate groups, comprised of more than 30,000 species with extreme variations in morphology and body plans [2]. For example, the ocean sunfishes (Molidae, Tetraodontiformes) exhibit one of the most impressive evolutionary transformations of the axial skeleton among ray-finned fishes [3,4]. It has been suggested that the enigmatic loss of caudal fin and caudal part of the vertebral column is related to “the existence in collections of a number of malformed adult tetraodontiforms without caudal fins” [5,6].

Various body and skeletal deformities in fish have been reviewed in many works [7-13]; earlier reports can also be found in Dawson’s bibliographies [13-17]. Many of these deformities apparently can be treated as monstrosities, particularly when the vertebral column is shortened without pronounced curvature, resulting in an altered body form that is extremely short and high. Such deformities are known in at least 26 species of 15 families (S1 Table). Remarkably, all individuals of Aischgrunder Karpfen, the South German carp breed (extinct since 1956), exhibited such deformity [18,19].

The riverine adaptive radiations in the large African barbs of the genus *Labeobarbus* Rüppell 1835 (Cyprinidae) – the dominating fish group in the waters of the Ethiopian Highlands – have been studied by us for almost 30 years. *Labeobarbus* belongs to the African Torini, a lineage of hexaploids [20-24] that originated in the Middle East via hybridization of tetraploids (maternal *Tor* lineage) and diploids (paternal *Cyprinion* lineage), and then dispersed throughout Africa [25]. Although the continental-scale phylogeny of *Labeobarbus* is still poorly resolved [26], the mt-DNA based phylogeny of this group in Ethiopian waters is relatively well-studied [27-32]. It is important to note that the barbs inhabiting the southeastern part of Ethiopia (drained by the Wabi-Shebele and Juba river systems into the Indian Ocean) form a monophyletic group, *Labeobarbus gananensis* (Vinciguerra, 1895) complex. This is a sister group to *Labeobarbus* inhabiting enclosed basins of the Ethiopian Rift Valley, as well as all *Labeobarbus* from the waters of western and northern Ethiopia belonging to the Omo-Turkana and Nile systems, and additional some Kenyan barbs [27,31,32].

In addition to the famed *Labeobarbus* radiation in Lake Tana [33-63], this group has also experienced parallel riverine radiations in four Ethiopian basins. These riverine radiations are similar in terms of their morphological and ecological differentiation, but differ in the ways of genetic divergence [32]. Outside of Ethiopia, the riverine *Labeobarbus* radiation seems to occur in the Inkisi River basin (Lower Congo) [64].

The *Labeobarbus* assemblage in the Genale River (the main tributary of the Juba River) is the only radiation occurring in southeastern Ethiopia. It is geographically separated from the other Ethiopian *Labeobarbus* radiations by the Ethiopian Rift valley. It exhibits the deepest divergence in terms of morphology, ecology and genetics compared to the other riverine radiations [32]. During an intensive sampling of the middle Genale River in 2009, we observed a substantial portion of barbs with extremely short and high bodies [31].

In 1993, an assemblage of the sympatric morphologically distinct morphs of the *L. gananensis* complex was discovered in the middle reaches of the Genale River [65]. This area was re-sampled twice in the 1990s [66,67], then in 2009 [27,31], and again in 2019. Together with the omnivorous generalized morph of *L. gananensis* and the scraping-feeder *L. jubae* (Banister, 1984), three additional morphs specialized in their morphology and feeding habits were always present in these samples from the middle Genale. These include the lipped morph of *L. gananensis* (having hypertrophied lips), the scraping morph *Labeobarbus* sp. 1 (provisionally called ‘smiling’) and the large-mouthed morph *Labeobarbus* sp. 2 (called ‘piscivorous’), as well as a hybrid morph called ‘smiling hybrids’. In 2009, a new morph, ‘short’, was discovered, which has an abnormally short and high body (Fig. 1). It composed approximately 10% of the total barb catch in both 2009 and 2019 samplings [31].

**Fig. 1.**
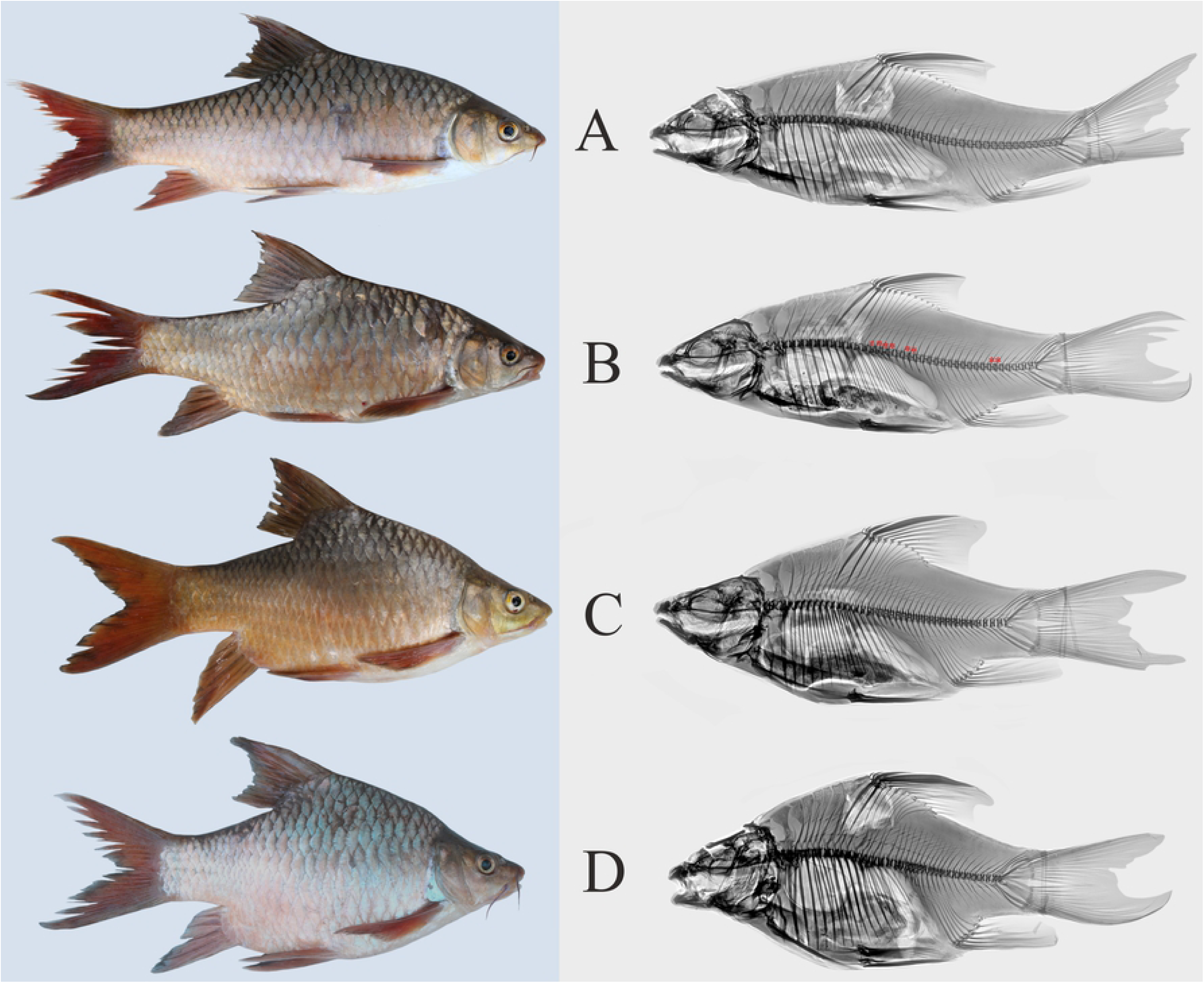
External appearance (left) and x-ray images (right) of the generalized and short *Labeobarbus* morphs from the middle Genale assemblage. A: generalized (SL = 242 mm), B: short, with eight deformed vertebrae, marked by red asterisks (SL = 245 mm), C: short, with 28 deformed vertebrae (SL = 216 mm), and D: short, with 31 deformed vertebrae (SL = 201 mm).

Trophic resource partitioning has been demonstrated among most morphs from the middle Genale *Labeobarbus* assemblage [31]. In contrast to the other morphs, the short morph does not diverge from the generalized morph in its diet [31]. Moreover, based on mtDNA markers, the short morph shares a common gene pool with the generalized and lipped morphs [31]. Preliminary investigation of the vertebral column revealed that each short individual had a substantial number of deformed (compressed and/or fused) vertebrae (Fig. 1).

The objectives of this study were: (1) to investigate the structure of the vertebral column in the short morph compared to other sympatric *Labeobarbus* morphs from the Genale River assemblage, (2) to analyze morphological differences between the short and related (generalized and lipped) morphs, with clarification of the contribution of the vertebral column shortening to these differences, and (3) to discuss the possible causes of the emergence of the aberrant morph in light of its life history characteristics and decade-long presence in the local *Labeobarbus* population.

## MATERIALS AND METHODS

### Ethics statement

Fish were collected in southeastern Ethiopia under the umbrella of the Joint Ethiopian-Russian Biological Expedition (JERBE), with permission from the National Fishery and Aquatic Life Research Center under the Ethiopian Institute of Agricultural Research and the Ethiopian Ministry of Innovation and Technology. Fish were sacrificed humanely using an anesthetic overdose (American Veterinary Medical Association). The experiments were carried out in accordance with the rules of the Papanin Institute of Biology of Inland Waters (IBIW), Russian Academy of Sciences, and approved by IBIW’s Ethics Committee.

### Study sites and sampling

The Ethiopian Highlands are divided by the Rift Valley into the western and eastern plateaus [68]. The southern part of the eastern plateau is drained by the Wabi Shebeli (Webi Shabeelle or Uebi Scebeli) and Juba (Jubba) river systems into the Indian Ocean via Somalia. Three large tributaries, the Genale (Ganale Doria), the Dawa (Daua) and the Weyb (Gestro), meet near the Ethiopia-Somalia border to form what is known as the Juba River within Somalia. The Genale, Weyb and Wabi Shebeli originate in the Bale Mountains, the highest part of the eastern plateau (Fig. 2).

**Fig. 2.**
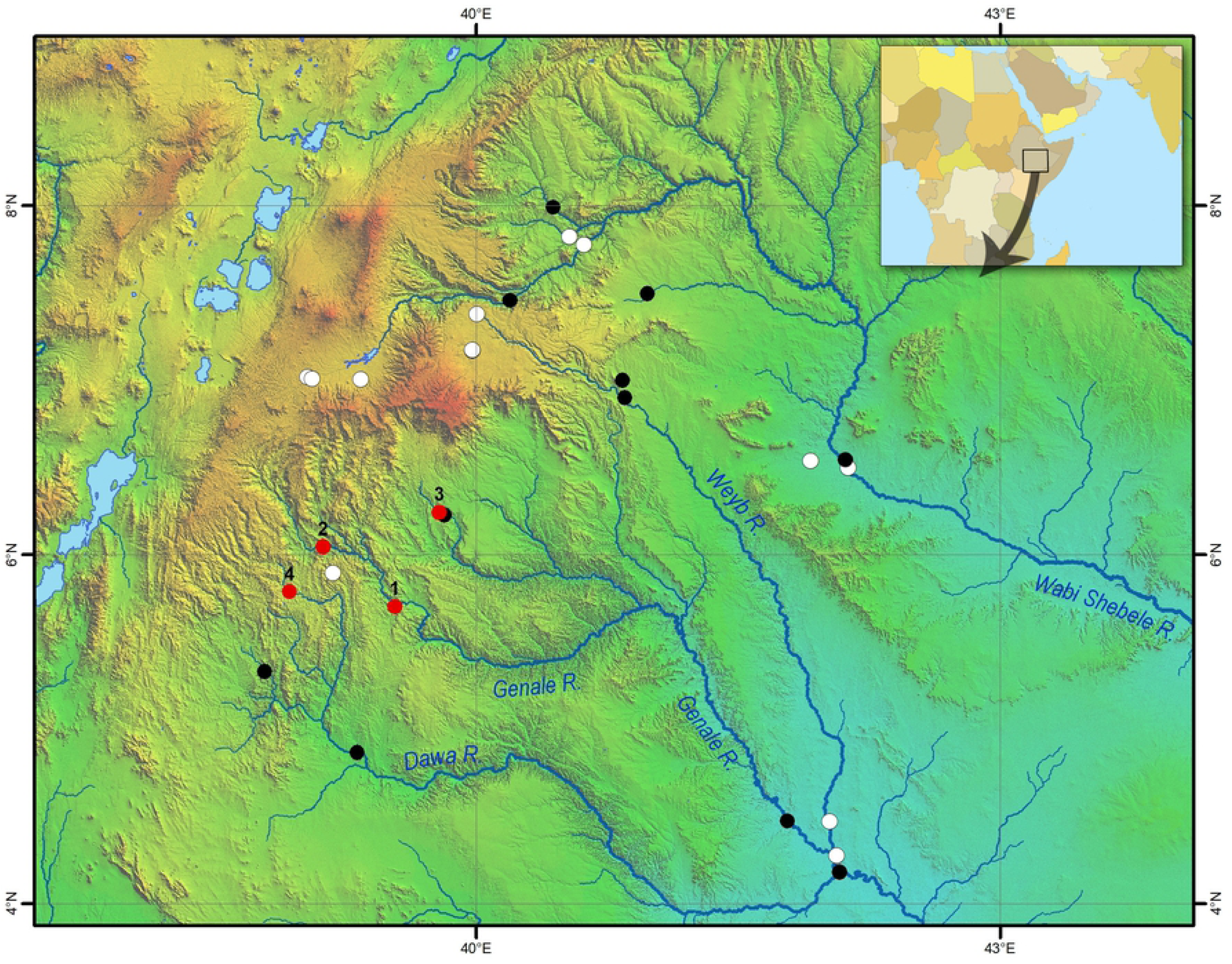
Map of sampling sites in the Juba and Wabi Shebele drainages. Black and red circles indicate sites where *Labeobarbus* was detected; white circles indicate sites where barbs were not found. Numbered red circles designate the most important sites for the current study – 1: Genale River, middle reaches, 2: Genale River, upper reaches, 3: Welmel River, and 4: Awata River. Geographical coordinates for all sites are given in S2 Table.

Since 1990, we have sampled a total of 28 sites across the Wabi-Shebele and Juba river systems (dots in Fig. 2). Among these, *Labeobarbus* were sampled from 15 sites, and not found in the other 13 sites. An additional four sites important for the present work are numbered in Fig. 2: (1) main sampling site, the middle reaches of the Genale River (5°42’ N 39°32’ E, altitude 1125 m above sea level, asl); (2) the upper reaches the Genale River north of Kibre Mengist (6°02’40” N 39°07’19” E, 1324 m asl); (3) the middle reaches of the Welemele River, a northern tributary of the Genale (6°14’30” N 39°47’22” E, altitude *c*. 1040 m asl); and (4) the upper reaches the Awata River, northern tributary of the Dawa (5°47’07”N 38°55’46”E, 1630 m asl). All sampling sites are listed in S2 Table. Most samples were taken in the different years from the end of January to mid-May during the main rainy season in southeastern Ethiopia [69], but before substantial rise of water level in the rivers.

The main sampling site (no. 1, Fig. 2) included two habitats: the continuous pool starting from the shallows around the island, characterized by moderate current and sandy/silty bottom, and the upper part of continuous rapids with small waterfalls, rocky bottom and very fast current (S3 Fig.). Samples were mostly taken from the pool by gill netting (at dawn and overnight), but an additional sample was obtained from the rapids with cast nets in 2009. The cast net catches differed substantially from the gill net catches, in terms of fish size and frequency of *Labeobarbus* morphs (S4 Table), therefore these catches were analyzed separately. The remaining sites (nos. 2-4) were sampled by both cast and gill netting during the day, and the catches were analyzed together.

The selected fish were maintained for several hours in large barrels submerged into the river, then killed with an overdose of MS-222 and preliminarily investigated for morphology. Tissue samples were taken for genetic and stable isotope analyses because the genetic and trophic specialization studies have been conducted on the same individuals [27,31,32]. Most specimens were preserved in 10% formalin in the field, and subsequently transferred to 70% ethanol in the laboratory. Of the 291 specimens collected in 2009, 132 were preserved with salt to make the dried bone preparations. These specimens were dissected for determination of sex and gonad maturity stages.

### Examination of vertebral column

Most formalin-preserved specimens were x-rayed. Based on the film or digital radiograms, the structure of vertebrae were examined. The total vertebral number and numbers of pre-dorsal, pre-anal, trunk, transitional and caudal vertebrae were counted according to Naseka [70]. Examination of vertebral structure and counts of vertebrae mentioned above were made by means of visual inspection of the dried bone preparations. The head magnifier (×4) was used for film and preparation inspection if necessary. In total, the vertebral column structure was studied in 364 Genale barbs representing all morphs. This number includes the radiograms of 34 and 45 specimens collected before 2009 and in 2019, respectively, as well as 285 radiograms and dried bone preparations of fish collected in 2009.

Witten et al. [11] propose a classification of vertebral column deformities that are repetitively observed in salmon (*Salmon salar* L.) under farming conditions. In our material, we observed all nine types of deformities united by these authors into the category of ‘compressed and/or fused vertebral bodies’. Taking into account the relatively small size of our samples, we did not consider the different types of such deformities separately. All vertebrae with compressed and/or fused bodies were considered herein as deformed vertebrae. The deformities of other categories [11] were not recorded in our material, except in one specimen of the smiling morph that had vertebral column curvature (kyphosis). Other deformities such as a presence of abnormal additional ribs and neural spines, and splitting of neural spine [71] were recorded in our material, however they were not analyzed because they were not related to the shortening of the vertebral column.

### Examination of other morphologic characters

Prior to preservation, the following measurements were taken (with a ruler, to the nearest 0.5 mm) on all specimens: standard length (*SL*), head length (*HL*), dorsal spine length (*DL*), maximum (*H*) and minimum (*h*) height of body, and length of caudal peduncle (*cpd*). For further analysis, the body length (*BL*) was calculated as *SL* minus *HL*.

Of all morphs sampled in 2009 (*N* = 129, *SL* = 98-308 mm), we randomly selected a subset of formalin-preserved specimens and took 23 measurements in the laboratory (with a caliper, to the nearest 0.1 mm): standard length (*SL*), head length (*HL*), snout length (*R*), orbit horizontal diameter (*O*), interorbital width (*IO*), postorbital length of head (*PO*), height of head at mid-orbit (*Ch*), height of head at occiput (*CH*), mouth width (*MW*), dorsal spine length (*DL*), pre-dorsal distance (*PrD*), post-dorsal distance (*PD*), caudal peduncle length (*cpd*), length of dorsal fin base (*lD*), maximum (*H*) and minimum (*h*) height of body, length of pectoral (*lP*) and ventral (*lV*) fins, pectoral-ventral distance (*PV*), ventral-anal distance (*VA*), height of anal fin (*hA*), length of anterior barbel (*Ab*), and length of posterior barbel (*Pb*). In the present work, only data on these extended measurements were included for 22 and 28 specimens of the generalized (*SL* = 108-257 mm) and short (*SL* = 119-244 mm) morphs, respectively. All measurements were taken by the same person for consistency [72].

In most sampled specimens, the following counts were taken: number of scales in the lateral line, number of scale rows above and below lateral line, number of predorsal scales, number of scales around the caudal peduncle, number of gill rakers on the first gill arch, number of branched rays in the dorsal fin.

### Examination of age and growth

In most salt-preserved specimens of the different morphs, age was determined from vertebrae. Annual rings were counted on the anterior and posterior sides of the fifth vertebra after removing the dried intervertebral discs. Counts were made with a binocular microscope at magnification ×16, with a drop of glycerin for contrasting. Age estimates were calibrated with vertebrae from the artificially reared *Labeobarbus* from Lake Tana [63].

Growth rate estimates for the different *Labeobarbus* morphs were based on comparisons of individual size variation in the different year classes. We did not calculate values of growth rate or parameters of the von-Bertalanffy equation because of the small sample sizes.

### Statistical analysis

Various R packages [73] realized in R-studio v.1.2.5033 were used for statistical analyses and plot construction: *summarytools* library was used for obtaining basal descriptive statistics, *ggpubr* library was used to analyze Pearson correlation and plot the result, *posthoc*.*kruskal*.*dunn*.*test* function was applied for posthoc Dunn’s test, *FSA* library was applied for Mann-Whitney U test, *prcomp* function was used for the principal component analysis (PCA) and plotting the results, *ggplot2* library was applied for plotting boxplots and histogram bars. The proportions of head and body were used for analyses of single characters, as well as for PCA – all measurements were divided by head length. Data was standardized for PCA.

### Deposition of material

All samples were deposited to the Severtsov Institute of Ecology and Evolution Russian Academy of Sciences and Papanin Institute for Biology of Inland Waters Russian Academy of Sciences under provisional labels of the Joint Ethiopian-Russian Biological Expedition.

### Abbreviation of morph names

The names of the morphs from the Genale *Labeobarbus* assemblage have been assigned the following abbreviations: short SH, generalized GN, lipped LP, *L. jubae* JB, smiling SM, smiling hybrids HB, and piscivorous PS.

## RESULTS

### Short definition

Initially, in the field, the short morph was identified based on altered body proportions: these individuals had relatively higher and shorter bodies compared to the normal individuals (Fig. 1). Preliminary investigation of the vertebral column revealed that each short individual had a substantial number of deformed vertebrae. The data on the structure of the vertebral column for all morphs are presented in Table 1.

**Table 1.** Occurrence of individuals with various numbers of deformed vertebrae in the different morphs in the total sample (1993-2019) (N, number of individuals; %dv, percent of individuals with deformed vertebrae in a particular morph).

| Morph | N | %dv | Number of deformed vertebrae per individual |  |  |  |  |  |  |  |  |  |
| --- | --- | --- | --- | --- | --- | --- | --- | --- | --- | --- | --- | --- |
|  |  |  | 0 | 1 | 2 | 3 | 4 | 5 | 6 | 7 | 8 | ≥ 9 |
| Shorts (SH) | 52 | 100 |  |  |  |  |  |  |  | 1 | 2 | 49 |
| Generalized (GN) | 78 | 16.7 | 65 | 1 | 8 | 2 | 2 |  |  |  |  |  |
| Lip (LP) | 26 | 11.5 | 23 |  | 2 | 1 |  |  |  |  |  |  |
| Jubae (JB) | 78 | 12.8 | 68 | 1 | 7 | 1 |  |  | 1 |  |  |  |
| Smiling (SM) | 48 | 14.6 | 41 | 1 | 4 | 1 |  |  | 1 |  |  |  |
| Hybrid (HY) | 36 | 5.6 | 34 |  | 2 |  |  |  |  |  |  |  |
| Piscivorous<br>(PS) | 46 | 9.8 | 41 |  | 3 |  | 1 |  |  |  | 1 |  |

We defined the SH morph as individuals with seven or more deformed vertebrae. It is important to note that the mouth structure in the SH morph was similar to that in GN and LP morphs that had normal vertebral columns. These morphs – with either shortened or normal vertebral columns – differed in the degree of lip development (S5 Fig.).

There were almost no individuals with markedly shortened bodies among the trophically specialized morphs (JB, SM, HY, PS). The only PS individual with eight deformed vertebrae (Table 1) displayed noticeable body shortening (S6 Fig.).

### Temporal and spatial distribution of the short morph

In the samples taken before 2009, SH individuals were not discovered among the 34 radiographed individuals of the different morphs, or among the hundreds of superficially examined barbs in catches of 1993, 1997 and 1998. To estimate SH prevalence in the samples of 2009 and 2019, we analyzed the total gill net catches. The catch composition appeared to be quite similar in 2009 and 2019 (S7 Table). The ratios of morphs remained roughly stable. It is important to note that SH prevalence in the total *Labeobarbus* catches was almost unchanged: they represented 10% of the 400 barbs sampled in 2009, and 11% of the 179 barbs sampled in 2019.

We did not sample the SH morph from the upper reaches of the Genale River (sampling site no. 2, Fig. 2), where most other morphs common to the main sampling site were found. Moreover, the SH morph was not recorded from other sampling sites in the Wabi-Shebele and Juba river systems (Fig. 2, S2 Table) or from other parts of Ethiopia (sampling sites may be found in [32,74,75]). However, at sampling site no. 3, we found barbs that had shortened bodies but without deformed vertebrae (discussed below).

### Short vs generalized and lipped morphs: plastic characters

As expected, body height of the SH morph was positively correlated (Fig. 3A) with the number of deformed vertebrae, which in turn was negatively correlated with body length (Fig. 3B). In other morphs of the Genale *Labeobarbus* assemblage, no such correlations were found.

**Fig. 3.**
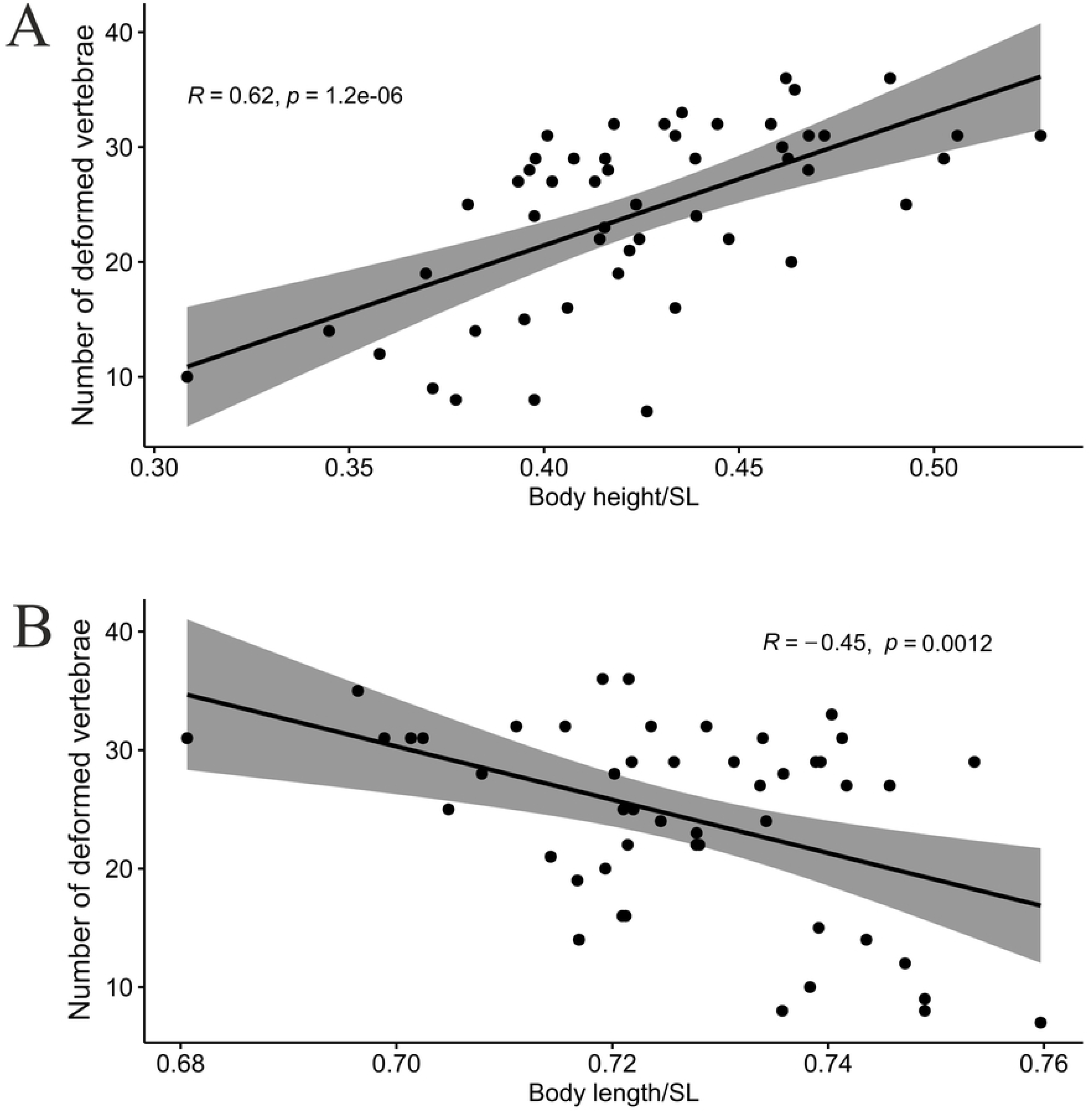
Pearson correlation of (A) body height and (B) body length with number of deformed vertebrae in the short (SH) morph.

The changes of body proportions in the SH morph (caused by the deformity of the vertebral column) resulted in a pronounced difference in appearance between this and the other morphs. Based on PCA, the SH and GN morphs were well differentiated from each other in PC1 (Fig. 4). Indices of 21 measurements relative to head length (*HL*) rather than *SL* were used in the PCA because of the variable influence of vertebral deformity on *SL* in the short individuals (Fig. 3B). PC1 explained > 45% of the variance, while PC2 was less than 24%. Eigenvectors of the 10 most loaded characters for PC1 and PC2 are given in S7 Table.

**Fig. 4.**
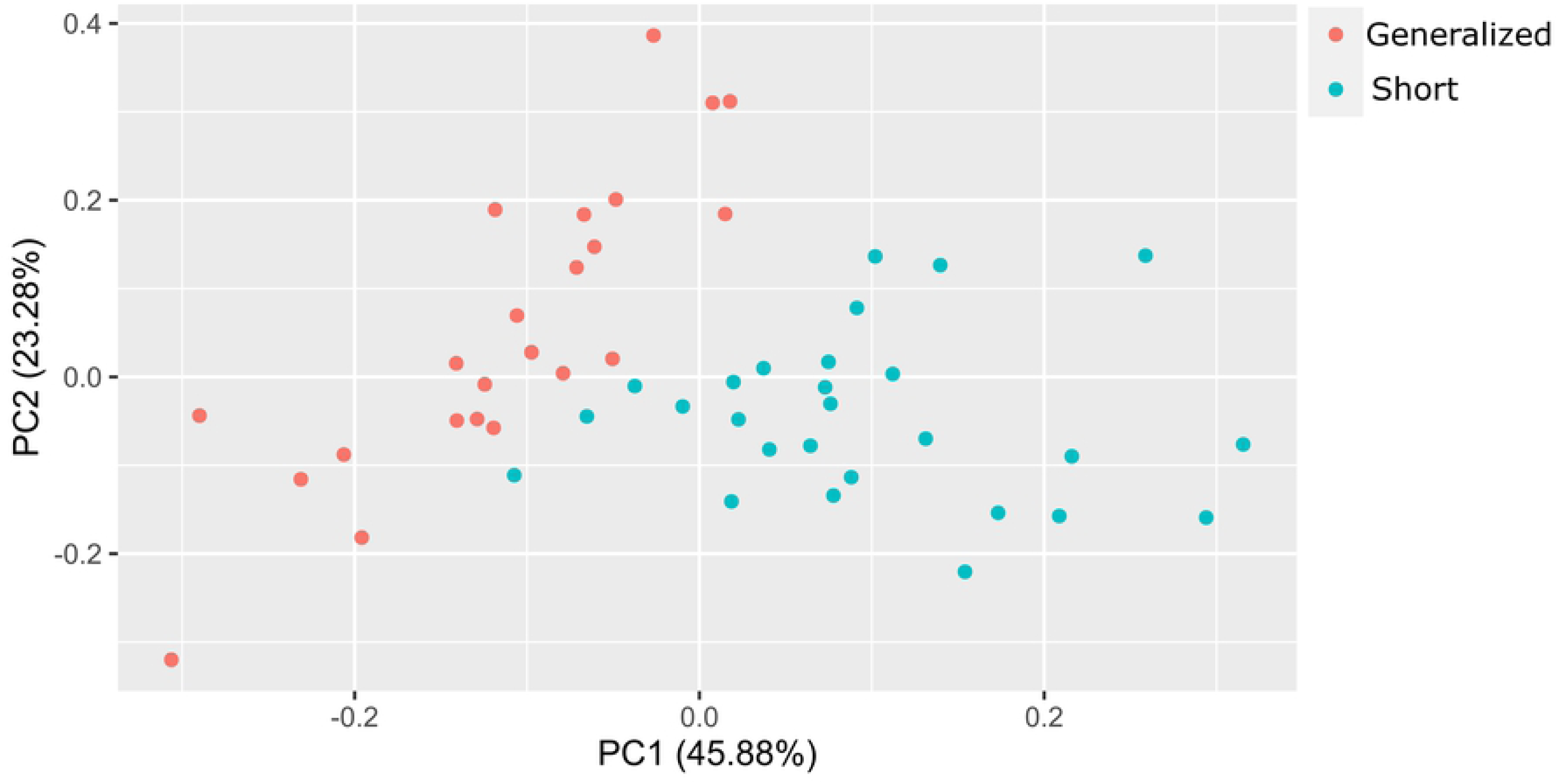
PCA of short (SH) and generalized (GN) morphs based on 21 indices of head and body proportions.

Using the a larger data set (which included the gill net catch of 2009, S5 Table), we compared the characters that had the strongest influence on body form (head length, heights of body and caudal peduncle) between the SH, GN and LP morphs (Fig. 5). The LP and SH morphs were similarly characterized by relatively longer heads (Fig. 5A). In the LP morph, the longer head length arose from elongation of the snout (S5 Fig.), while in the SH morph the head length appeared longer relative to the shortened body. The LP, SH and lipped SH morphs all had relatively longer heads than the GN; the lipped SH morph had a notably longer head than the SH morph, but the difference was not significant (Fig. 5A). The relative heights of body and caudal peduncle in the SH morph differed significantly from both GN and LP, however there were no such differences between the lipped SH and any of the other morphs (Fig. 5B-C). Notably, lipped SH exhibited the most variation in relative heights of body and caudal peduncle compared to the other morphs. This variation was caused by different degrees of snout elongation and variation caused by body shortening (determined by varying number of deformed vertebrae), which seemed to act additively.

**Fig. 5.**
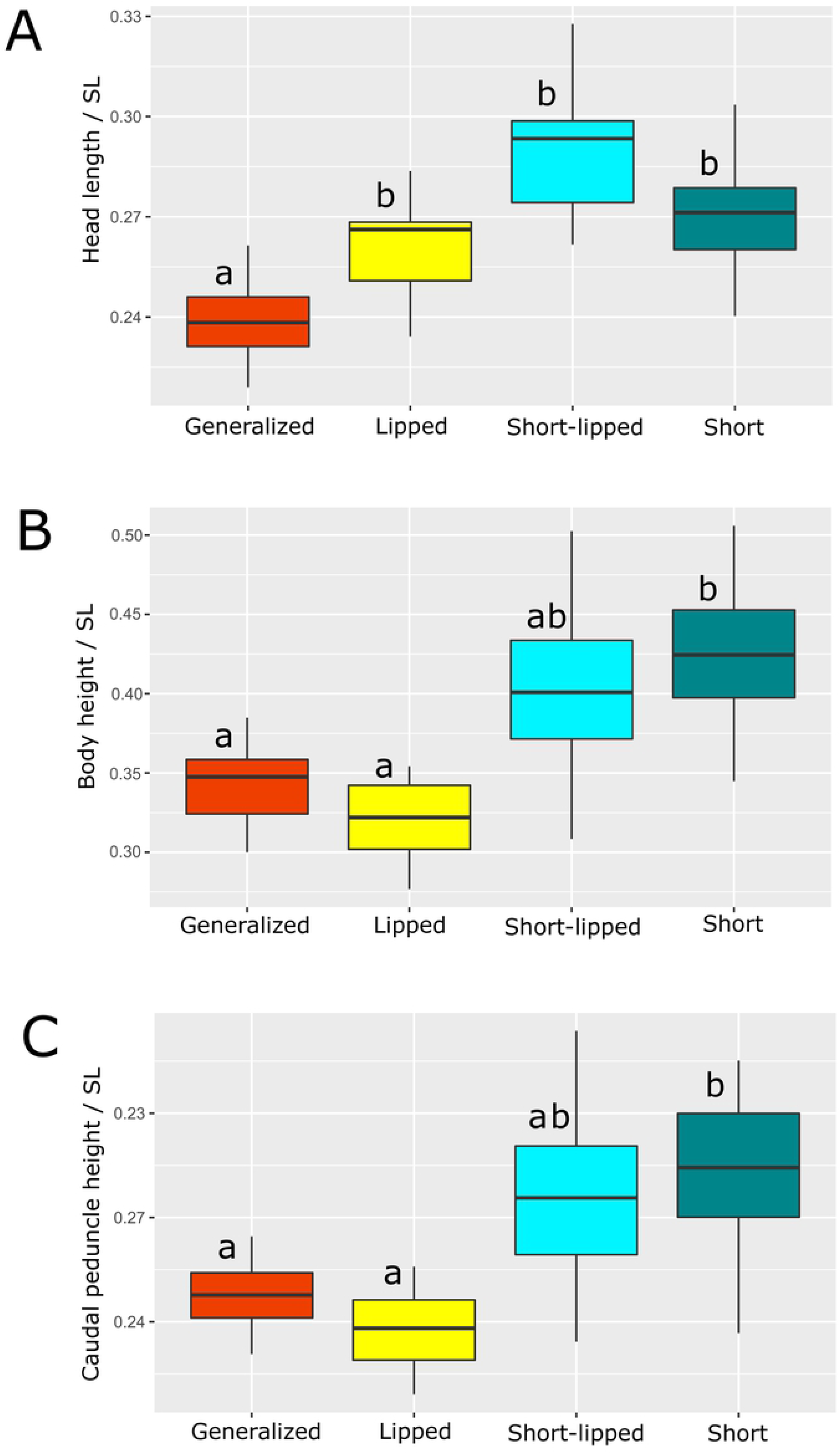
Indices of (A) head length, (B) body height, and (C) caudal peduncle height in generalized, lipped, lipped short, and short morphs sampled in 2009. Median is shown as the horizontal black line inside the box. The box represents 1st and 3rd quartiles of variation. Lowercase letters above the boxplots indicate significant differences between morphs (*p* < 0.05, Dunn’s test with Holm adjustment of *p*-value).

The comparison of barbs sampled in 2009 and 2019 yielded unexpected results. Although the later sample was rather small, significant differences in relative head length and height of caudal peduncle, as well as a nearly significant (p = 0.054) difference in body height, were revealed between GN sampled in 2009 and 2019 (Fig. 6). Remarkably, in only 10 years the GN morph became significantly more similar to the SH morph in all three of the indices most important for their discrimination. No differences in *SL* and absolute values of the corresponding measurements were found between samples of 2009 and 2019 (Mann-Whitney U test).

**Fig. 6.**
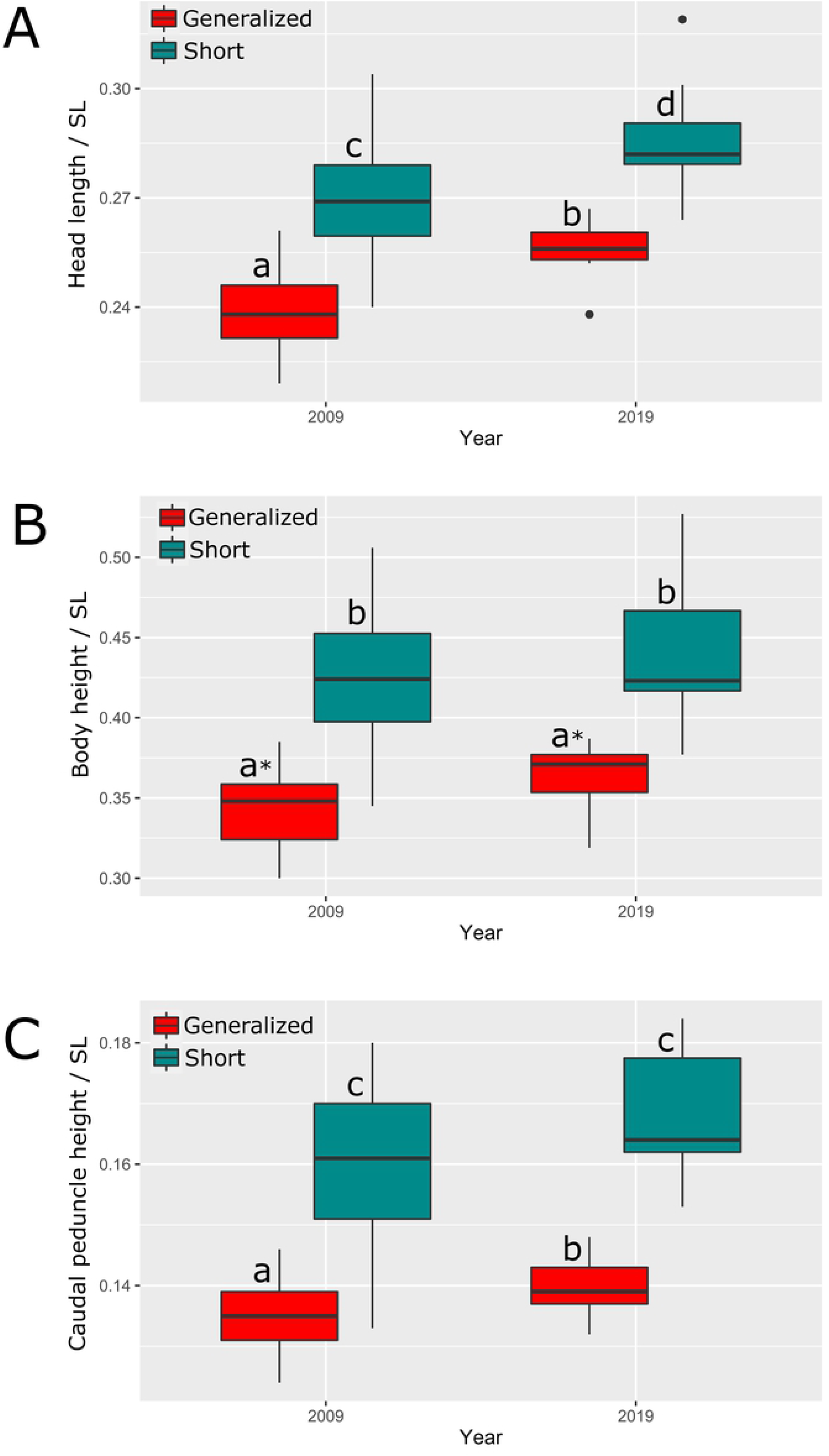
Indices of (A) head length, (B) body height, and (C) caudal peduncle height of generalized and short morphs sampled in 2009 and 2019. Median is shown as the horizontal black line inside the box. The box represents 1st and 3rd quartiles of variation. Lowercase letters above the boxplots indicate significant differences between morphs (*p* < 0.05, Mann-Whitney U test; * - nearly significant, *p* = 0.054).

### Short vs generalized and lipped morphs: structure of vertebral column and other counts

Comparison of vertebral counts between the SH morph and the GN and LP morphs did not reveal any differences in total number of vertebrae or in numbers of vertebrae in the three segments of the vertebral column (trunk, transitional and caudal). Moreover, there were no differences in the numbers of pre-dorsal and pre-anal vertebrae (S9 Table).

Distribution of the deformed vertebrae along the vertebral column was not random, and was different in the SH morph compared to the other morphs (Fig. 7). When the data on the positions of the deformed vertebra was merged for individuals with the different total number of vertebrae, the frequency estimates for each specific vertebra in the caudal region were obscured. For this reason, only the data for individuals with 41 total vertebrae are presented in Fig. 7. The respective data for individuals with 40, 42 and 43 vertebrae are shown in S10 Figure.

**Fig. 7.**
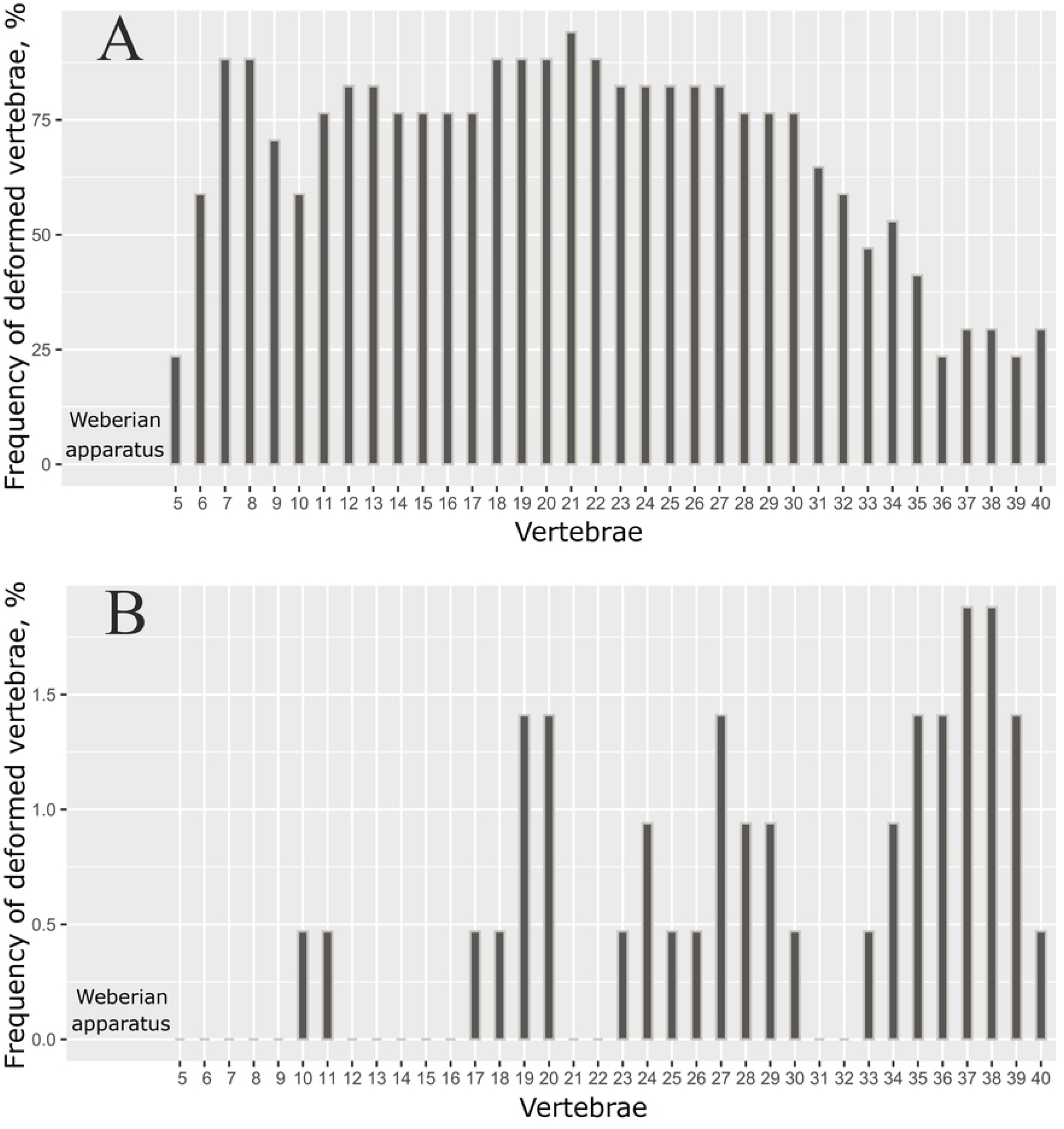
Distribution of deformed vertebrae along the vertebral column in (A) short morph (n = 17), and (B) other morphs (n = 47). Only data for individuals with 41 vertebrae are shown here; the deformities were not analyzed for the first four vertebrae (Weberian apparatus) or the 41^st^ (pre-ural 1) vertebra for technical reasons.

As shown in Fig. 7A, the lowest frequencies of each specific vertebra deformity in the SH morph were found in 5th vertebra, and also in the caudal region, where the minimum frequencies were found in the 36-40^th^vertebrae. In the other morphs, however, the deformed vertebrae occurred mostly in the posterior half of the vertebral column, with maximum frequencies in the 37-38^th^ vertebrae (Fig. 7B).

No differences were found in the counts of scales, gill rakers and fin rays between the GN and LP morphs, on the one hand, and the SH and lipped SH morphs, on the other hand. No changes in the counts for samples 2009 and 2019 were recorded.

### Short vs generalized and lipped morphs: size, age, growth rate, sex ratio and gonad conditions

The range of size variation in the SH morph (*SL* 107-291 mm) was less than that in any other morph from the middle Genale assemblage in 2009 and 2019. The ranges for the GN and LP morphs were 51-375 mm and 106-459 mm, respectively (S4 Table). Most of the salt-preserved barbs were aged from vertebrae. A maximum age of six years was recorded for the SH and HB morphs, seven years for the SM morph, eight years for the GN morph, nine years for the JB morph, 12 years for the LP morph and 14 years for the PS morph. There was no correlation between the percentage of deformed vertebrae and age.

Analysis of size variation within each year class of the SH and GN morphs revealed that when *SL* was used as a measure of linear growth, SH did not differ in size from GN during the first four years (S11 Fig.). Modification of the vertebral column in the SH morph, however, apparently resulted in shortening of their body and respective reduction of *SL*. Hence, it is reasonable to consider the head length as a measure of linear growth in comparisons of the SH morph with the other morphs. In this case, the SH morph demonstrated a markedly fast growth rate during the first four years, which then leveled off during the fifth and sixth years (Fig. 8).

**Fig. 8.**
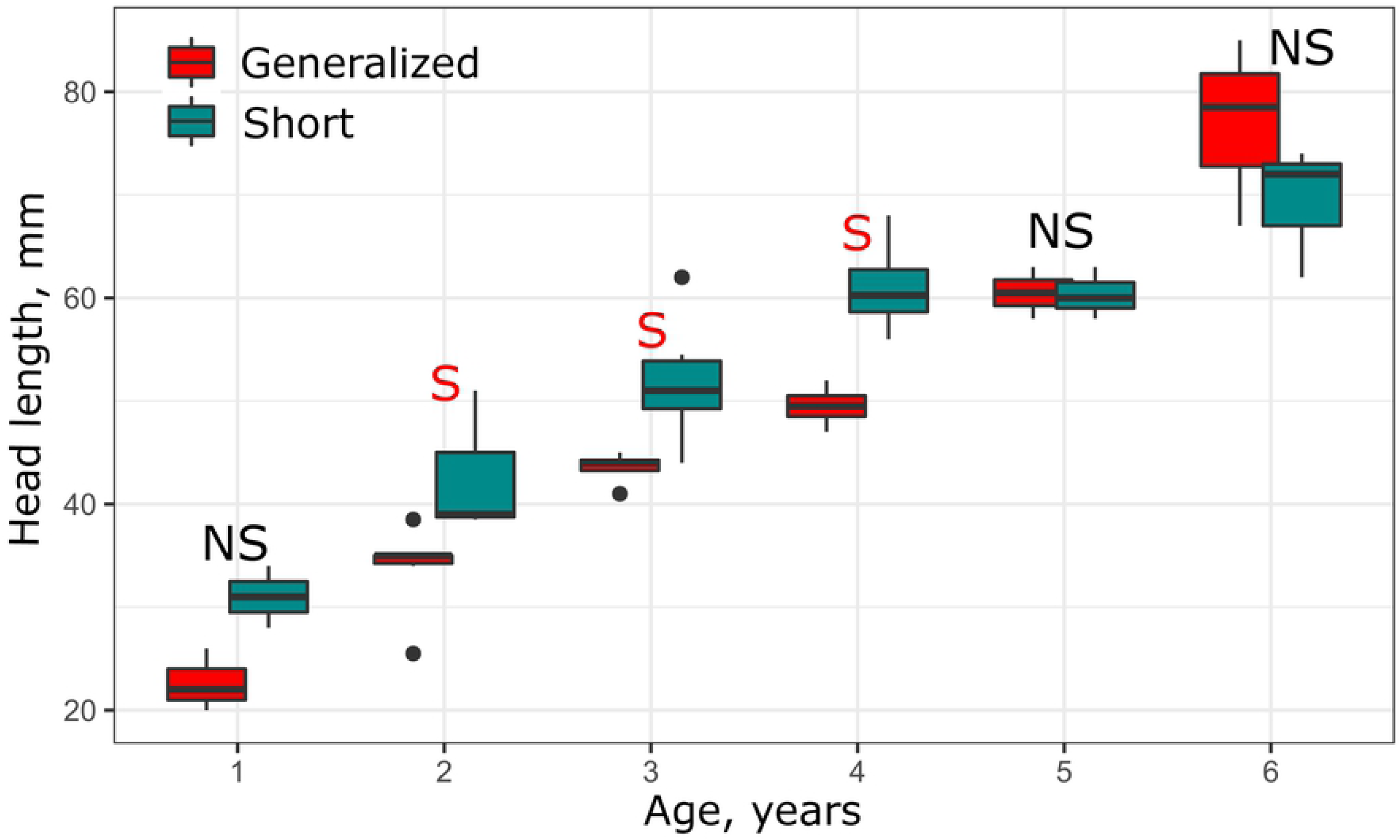
Age *vs* head length in generalized (GN) and short (SH) morphs in 2009. Median is shown as the horizontal black line inside the box. The box represents 1st and 3rd quartiles of variation. Letters above each boxplot designate significant (red S), or non-significant (black NS) differences between morphs within each year class, based on Mann-Whitney U tests (*p* < 0.05).

Gonads were examined in 30 SH individuals: five juveniles, 11 females and 14 males were recorded. There were 16 fish at maturity stage I, 11 fish at stage II, three fish at stages II-III and III. Among the 46 examined individuals of GN and LP morphs, 16 juveniles, 14 females and 16 males were recorded. There were 24 fish at maturity stage I, 20 fish at stage II, one fish at stages II-III and one female at stage IV. Thus, neither sex ratio nor gonad condition differed in the SH morph compared to GN and LP morphs.

## Discussion

It is impossible to clearly distinguish the SH morph from the GN and LP morphs using only the plastic characters reflecting their body form, which aligns with Darwin’s observation that “monstrosities … graduate into varieties” [1]. Discrimination is possible, however, by using the number of deformed vertebrae per individual: the GN and LP morphs have ≤ 4 such vertebrae, while the SH morph has ≥ 7 (Table 1). Considering the emergence of the SH morph in the *Labeobarbus* population predominantly composed of GN and LP morphs, one may suggest that it is determined by some factor(s) acting qualitatively or the threshold reaction of the *Labeobarbus* individuals to some factor(s) with continuously varying parameters.

In regards to Darwin’s idea about the propagation of monstrosities “for a succession of generations in a state of nature” [1], we are motivated to estimate the period of SH existence in a population. Taking into account the absence of SH individuals in our sample from 1998 and their maximum age of six years in the sample from 2009, it is reasonable to suggest that the SH morph appeared in the population between 1999 and 2005. The minimum size of SH individuals in the sample from 2019 (*SL* = 177 mm) corresponds to an age of 2-3 years according to the SH sample from 2009 (S4 Table). Thus, the SH individuals regularly appeared in the population from 2005 (or slightly earlier) until at least 2016. This situation could be explained by the influence of some external factor(s) that abruptly appeared and then became permanent in the environment, or by the reproduction of SH individuals during the ten year period, which obviously includes the life-span of several generations.

A central question has arisen from the discovery of the SH morph: does the emergence and 10-year persistence of this morph have a genetic background? Based on mtDNA variation, there is no significant genetic difference between the SH, GN and LP morphs [31], but it is still possible that these differences are present but subtle. In the absence of direct genetic evidence, the factors causing vertebral deformities in fish are considered more generally.

### Factors responsible for abnormal body shortening in fish

There is extensive literature devoted to the question of vertebral deformities in fish. The most comprehensive data available pertains to artificially propagated fish species, especially farmed salmonids. Fish with abnormally shortened bodies displaying multiple compressed and/or fused vertebrae are quite frequent in farmed stocks of Atlantic salmon *Salmo salar* L. 1758 [11,76-86]. Externally (imaged in [**76])**, they are similar to the short morph from the Genale *Labeobarbus* assemblage. These salmons, called ‘short tails’ or ‘short-spined’ individuals [78,81], pose a serious problem for the farming industry because of their reduced commercial value [87]. Therefore, special efforts have been undertaken to elucidate the factors determining the appearance of short tails, however the results are vague, at least in terms of the genetic factors. Initially, results suggested that the deformities were heritable [76,79], but this was questioned later [84,85]. As described by Witten et al. [81, p. 244]: “one cannot rule out the possibility that some fish are genetically predisposed and develop vertebral compression as a reaction to external cues”.

The factors that increase the risk of vertebral deformities in farmed Atlantic salmon and other salmonids are given by Witten et al. [11]: bacterial and parasitic infections, vitamin C deficiency, phosphorous deficiency, elevated egg incubation temperature, fast growth in under-yearling smolts, inappropriate light regimes, vaccination, inappropriate water current and quality, as well as environmental pollution. All of these factors (except vaccination), along with radiation [71] may be present in the natural population of the middle Genale *Labeobarbus*. Notably, there is a relationship between the increased frequency of vertebral deformities and high growth rate in Atlantic salmon smolts [11,82,85]. We also found an increased growth rate in the SH morph compared to the GN and LP morphs at the age of 2-4 years. Thus, the deformations of the vertebral column in the SH morph could be caused by the accelerated early individual growth and mechanical muscle overload of bones, as suggested for Atlantic salmon [11,82,85]. For the Atlantic salmon post-smolts, a retardation of growth is reported in fish with high numbers of deformed vertebrae [86], however the authors used fork length as the measure of linear growth, which may substantially bias the growth estimates.

As for other salmonids, a short tailed brown trout, *Salmo trutta* L. 1758 was reported from Scotland over a century ago [89]. Much later, >5% of the brown trout from a fish farm in Hampshire (England) exhibited the short tail phenotype and extensive compression of the vertebral column [90]. The author of the study indicated that inbreeding depression was a probable cause of the abnormalities. However, this explanation is hardly applicable to the Genale barbs for two reasons. First, the generalized morph, which is the ancestor of most individuals of the short morph, is the most common morph among the middle Genale barbs, and hence is unlikely to suffer from inbreeding depression. Second, spatial subdivision of the Genale barb populations can occur only in the spawning grounds; in this case, an increase of inbreeding would only result from almost perfect homing, which is highly unlikely in *Labeobarbus*.

In a study of the vertebral fusion patterns in coho salmon, *Oncorhynchus kisutch* (Walbaum 1792) [91], the author demonstrated that the distribution of vertebral fusions along the spinal column differed significantly among crosses from two hatchery stocks, indicating a genetic basis for this character. Recently, differences in the distribution of vertebral fusions along the spinal column were found between the genetically distinct year-classes of Atlantic salmon and its hybrids with brown trout and Arctic char *Salvelinus alpinus* (L. 1758) [92]. This result is interesting for understanding the difference in distribution of deformed vertebrae along the spinal column between the SH morph and other Genale *Labeobarbus* (Fig. 7).

In non-salmonid fishes, it has been reported that body shortening from the compression of the vertebral column was apparently heritable in the south German common carp Aischgrunder Karpfen [18], as well as in laboratory strains of the banded topminnow, *Fundulus cingulatus* Valenciennes [93] and two poeciliids, blackstripe livebearer *Poeciliopsis prolifica* Miller 1960 [94] and guppy *Poecilia reticulata* Peters 1859 [95]. In zebrafish, *Danio rerio* (Hamilton 1822), column shortening and vertebral fusions are exhibited by type I collagen mutants, as well as by individuals with knockout alleles in two genes involved in type I collagen processing [96]. At the same time, the increased frequency of vertebral deformities in wild-type zebrafish is induced by high rearing densities [97]. The stumpbody phenotypes with the shortened vertebral column in channel catfish *Ictalurus punctatus* (Rafinesque 1818) [98] and blue tilapia *Oreochromis aureus* (Steindachner 1864) [12,99] are described as not heritable.

The situation in the Japanese rice fish, *Oryzias latipes* (Temminck & Schlegel 1846) is especially informative [100-102]. The shortened vertebral column and vertebral fusions are found in (1) *fused* mutants obtained from the laboratory strain (‘a simple recessive Mendelian character’ with expression modified by temperature [100,102]), (2) ‘wild-fused’ phenotype from ‘a certain region on the eastern outskirts of the City of Nagoya’ that were not heritable, and (3) individuals of the normal strain treated by phenylthiourea at the early embryonic stage [101]. In general, it is obvious that in different species and even in different lineages of the same species, the emergence of fish with shortened columns and fused vertebrae may be determined by various external and genetic cues.

Considering the possible role of external cues, the following points must be mentioned. First, among the middle Genale *Labeobarbus* assemblage, the deformation of the vertebral column is found almost exclusively within the isolated genetic pool including the GN, LP, and SH morphs but excluding the trophically specialized morphs. If the deformation has no genetic background and is triggered by external cues such as mechanic, thermal, chemical or radioactive environmental stress, these impacts are only acting selectively. For example, they would have temporal or spatial effects on only some of the GN and LP barbs, but not the rest of the assemblage that is possible, in our understanding, mostly on the spawning grounds.

Second, environmental stress usually results in multiple malformations of different morphological structures [71,103]. We did not observe such multiple morphological abnormalities in SH. Third, external cues like bacterial [104] or parasitic [105,106] infection, and nutrient deficiency (*e*.*g*., vitamin C or phosphorous [107]) are usually manifested as a decrease in the individuals’ conditions, especially in the growth rate, which is often retarded in poor health. However, the growth rate was exceptionally high in the abnormal individuals (SH) observed here. Taking into account all of the above, we suggest that genetic factors (possibly together with some environmental ones) are the most parsimonious explanation of the emergence and 10-year persistence of the SH morph in the middle Genale *Labeobarbus* assemblage.

Nevertheless, to definitively prove this suggestion, we must seek answers to the following questions: do SH individuals reproduce in nature? Would be the progeny of artificial crossing be different for the SH breeders vs. normal GN and LP breeders? Are there any genetic/genomic differences between the SH morph and other morphs? Hopefully, these will be addressed in further studies.

### Shortened vertebral column as a target of natural selection

Among several streams in the State of Washington, Patten [108] noted that ‘the presence of many severely deformed fish (some of a mature size) shows that some factor or factors permitted their survival’. Möller [109] also stated that ‘anatomical abnormalities … may become particularly evident in populations that do not suffer from a high selection pressure by predators or food competitors’. Essentially these authors are referring to a relaxation of stabilizing selection; an influence of natural selection is often reasonable to expect, but very difficult to prove unambiguously. There are many records on the increased frequency and severity of skeletal abnormalities in artificially reared fish stocks compared to natural populations [110-117]. Interestingly, the opposite situations are also reported [117,118]. Demonstrating how the intensity of rearing conditions plays a role in increasing the frequency and severity of skeletal abnormalities for several fish species [117,119-122] can lend support to the role of selection relaxation, both in cultivated stocks and natural populations exhibiting the mass abnormalities.

Some researchers [123-125] have indicated the relaxation of natural selection as an important mechanism in increasing variation of morphological, ecological and other traits in the course of adaptive radiation. Such an increase often allows the radiating species to occupy new adaptive zones. This phenomenon has been called ‘extralimital specialization’ by Myers [126]. In this perspective, the occurrence of the vertebral deformations at a relatively high frequency only in the radiating *Labeobarbus* assemblage in the middle Genale River may be evidence for relaxed selection in this particular site.

Concerning the possibility of directional selection – positive or negative – in shaping the abnormal barbs (SH morph) in the middle Genale River, the following points must be mentioned. First, some retardation of growth in the 5-6 year classes of the abnormal barbs (Fig. 8), along with their relatively short life span (< 7 years), could be treated as evidence for negative selection at later stages of ontogenesis. For Atlantic cod, *Gadus morhua* L. 1758 from the German Wadden Sea, a few individuals with spinal compression were found in the larger size classes [127]. This reduced frequency in later life stages was explained by negative selection for this phenotype. At the same time, seasonal variation of spinal compression frequencies in the smaller size classes was explained by the migratory activity that differed between the abnormal and normal individuals [127], where the former are considered to be less active.

Second, one may expect that the barbs with deformed vertebral columns should exhibit reduction in swimming performance, but this is not straightforward. Studying the triploid lines of Atlantic salmon, Powell et al. [128] did not find a difference in swimming performance between normal fish and those with visible spinal shortening. They did note a poor recovery from exhaustive swimming exercises in the deformed fish compared to the normal fish. Based on this result, we cannot exclude the possibility of variable swimming performances between the deformed and normal Genale barbs. At the same time, it is clear that the deformed barbs can avoid habitats with fast water current in order to reduce the energy cost of their abnormality.

Third, there could be various reasons for the difference in distribution of the deformed vertebrae along the vertebral column between barbs with few and many compressed and/or fused vertebrae (Fig. 7). As described earlier, most of these vertebrae in the individuals with slightly deformed vertebral columns occur in the posterior end of the fish [9,83,86,97,121,129,130]. This phenomenon is explained by the morphogenetic influence of the caudal complex, whose normal development includes several vertebral fusions [130]. The increased frequency of abnormalities in the caudal region was also detected as a result of thyroid hormone disruption [131,132]. However, in the deformed (SH) barbs, the frequency of deformed vertebrae was lower in the pre-caudal region (Fig. 7). This is unlike the farmed stock of Atlantic salmon [86]. In the wild Genale *Labeobarbus*, this frequency distribution of deformed vertebrae may be caused by the genetic distinctiveness [91,92] of these deformed individuals, or by selection against the individuals with deformed vertebrae in the caudal region, for example because of their potentially decreased swimming capability.

The data obtained from individuals with normal vertebral columns allows us to shed light on the susceptibility of column deformations to directional natural selection. First, comparison of the samples from 2009 and 2019 unexpectedly revealed changes in the body form of the GN morph (Fig. 6). The GN individuals became more similar to the SH morph, particularly in the relative body height. At the same time, the SH morph also became more high-bodied than in 2009. The most parsimonious explanation for this phenomenon is a selective pressure for the increase in relative body height in the population studied. Second, in our small sample from the Welemele River (the northern tributary of the Genale; Fig. 2, site no. 3), together with the normal generalized *Labeobarbus* morph, we found a few individuals with body proportions very similar to that in the SH morph from the middle Genale assemblage. However, their vertebral columns were not deformed (Fig. 9). This finding might provide evidence for a selective pressure for an increase in the relative body height in another *Labeobarbus* population.

**Fig. 9.**
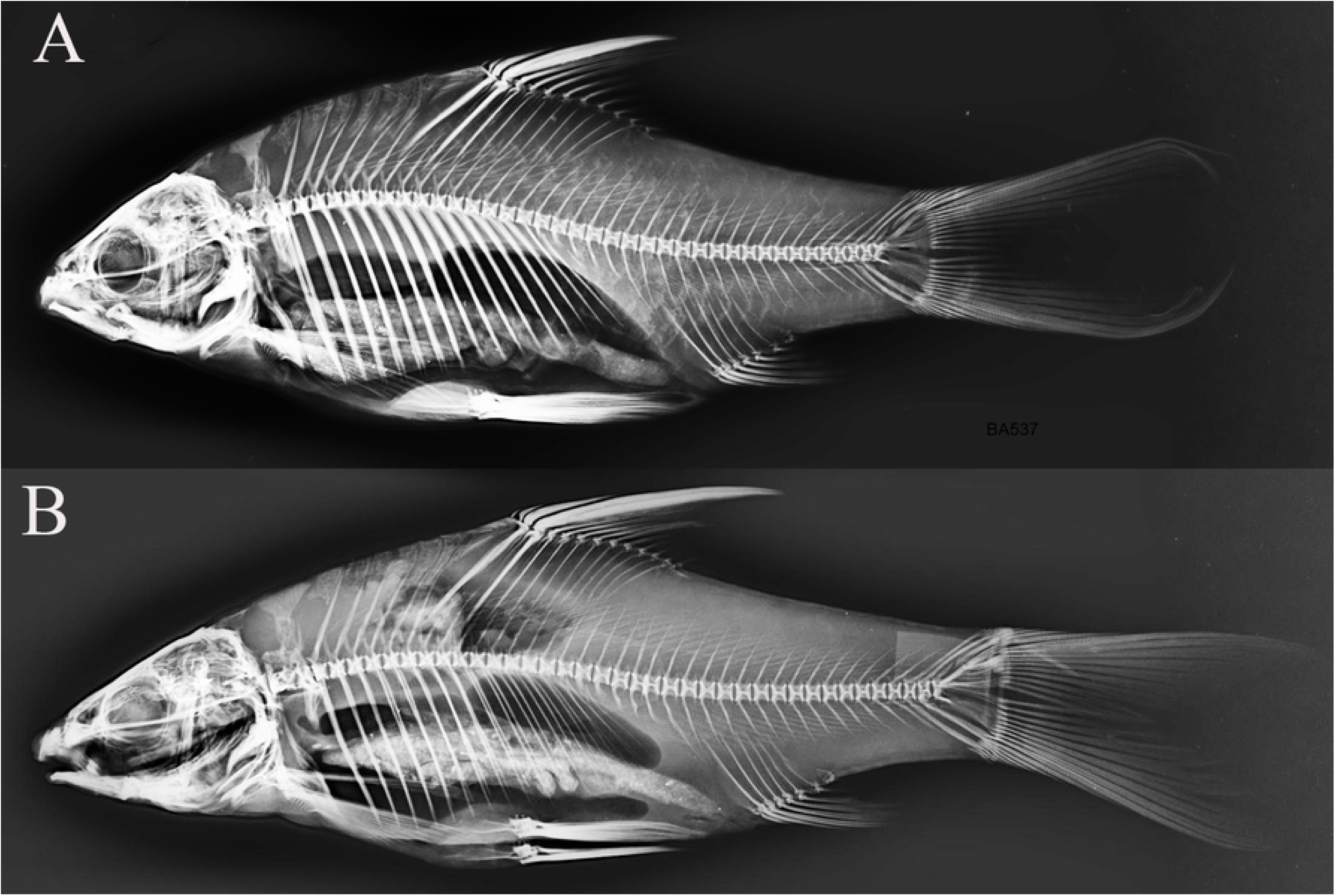
X-ray images of two *Labeobarbus* from the Welemele River with (A) short-like exterior (*SL* = 160 mm) and (B) normal exterior (*SL* = 177 mm). No deformed vertebrae were detected in either individual.

The increased body height is considered as an anti-predator adaptation in the crucian carp *Carassius carassius* (L. 1758) [133,134]. Indeed, the differentiation among *Labeobarbus* in the Chamo-Abaya lake basin (Ethiopian Rift valley) corroborates this idea. The lakes are inhabited by the high-bodied barbs classified as *L. bynni* (Fabricius 1775) [135], while the tributaries are inhabited by the low-bodied barbs classified as *L. intermedius* (Rüppell 1835) [135]. However, phylogeny based on mt-DNA [30] demonstrates a close relationship between these two forms and their distant relation to *L. bynni* from the Nile basin. Thus, high-bodied forms evolved independently in the Chamo-Abaya and Nile basins most probably as an anti-predator adaptation, as the faunal data [74,136,137] indicate the greater predator diversity in these basins compared to the Chamo-Abaya tributaries and most other rivers of the Ethiopian Highlands that are populated by the low-bodied *Labeobarbus*. In our understanding, the importance of the predator-induced high-bodied phenotype is evident in some *Labeobarbus* populations. In these cases, natural selection could favor the Genale deformed barbs (SH morph) because of their anti-predator high-bodied phenotype.

### Shortened vertebral column as a population phenomenon

In natural fish populations, the frequency of individuals with few compressed and/or fused vertebrae usually ranges from less than one percent to several percents (usually <10%) [109-112, 114,118,138-143], but sometimes it appears substantially higher (up to or over 50%) [115,117,143-145]. Considering these facts, the frequencies of individuals with 1 to 6 compressed and/or fused vertebrae (ranging from 5% to 15% in the morphs other than SH; Table 1) do not look outstanding. However, among the 34 individuals sampled from the Awata River (Fig. 2, site no. 4), such deformed vertebrae were not found at all. Hence, in the absence of data for other *Labeobarbus* populations, we may consider the frequency of individuals with deformed vertebrae in the middle Genale populations somewhat increased. Unfortunately, we cannot say with certainty how this is related to an emergence of the SH morph there.

There are many reports of individuals with the markedly shortened bodies and deformed vertebral columns in natural populations of different fish groups (S1 Table). Among wild cyprinids, there are reports of a Mesopotamian barb, *Mesopotamichthys sharpeyi* (Günther 1874), with nine deformed vertebrae, and a yellowfin barbell, *Luciobarbus xanthopterus* Heckel 1843, with 22 deformed vertebrae [146,147]. The situations, as in the Genale *Labeobarbus*, where the abnormality becomes a population phenomenon occurring with substantial frequency among several successive generations are rare. It is important to note that in the Genale barb assemblage, only two morphs (GN and JB) were always present in catches with frequencies higher than 10% recorded for the abnormal (SH) morph (S7 Table).

To the best of our knowledge, apart from the Genale *Labeobarbus*, the similar situations are described for only two other fish species, Atlantic cod *Gadus morhua* from the Elbe estuary and German Wadden Sea [127,148-150] and Arctic char *Salvelinus alpinus* from Transbaikalian Lake Dzhelo [151]. The high prevalence of vertebral column deformities was recently reported in a wild population of lumpfish *Cyclopterus lumpus* L. 1758 from Masfjorden, Norway [117], but changes of the body form and temporal stability of the abnormal phenotypes are not evident in the latter species.

In 1971, Wunder [148] reported the high prevalence (10-15%, sometimes up to 20%) of cod individuals with shortened bodies and deformed vertebrae in catches from the Elbe estuary and adjacent regions of the Northern Sea. In each deformed individual, 40-80% of vertebrae were compressed and/or fused. This abnormality was most frequent in the young cods (length < 20 cm). The situation persisted at least until the late 1980s [127,149,150]. The geographical range of this cod deformity expanded to some regions of the Wadden Sea; the mean frequency of the deformed individuals in the catches of all inshore regions of the German Wadden Sea was 9.2% in the size class of 11-23 cm [127].

Dzhelo is a small lake (1.4×0.3 km, maximum depth 25 m) in the upper reaches of the Lena River system in Transbaikalia. According to Alekseyev [151], the lake is inhabited by dwarf char (maximum length 24 cm). There are numerous deformed individuals with the shortened body, which also show multiple fusions of vertebrae. Their prevalence is 12%, and a change of the body form in the individuals with ≤ 6 deformed vertebrae is not noticeable. The lake was sampled once in 2003 [151].

The main question about these deformities in cod and char is whether they still persist with the substantial frequencies. Comparative etiology of vertebral deformities in cyprinid, salmonid and gadid species could be of great interest. Moreover, deeper understanding of the action of natural selection on these abnormalities can shed the light on how morphological novelties are established in a population, especially if the abnormalities in question have the genetic background.

## Conclusions

The striking emergence of individuals with deformed (shortened) vertebral columns resulting in a high-bodied phenotype is a phenomenon rarely detected among wild fish populations. For the particular case of the radiating *Labeobarbus* assemblage from the middle Genale River, the following circumstances must be highlighted: i) the recent emergence (∼ 15 years ago) of the high-bodied phenotype with the shortened vertebral column, ii) the persistence of this phenotype with a substantial frequency (∼ 10%) in several generations (>10 years) in a population, iii) the fact that this is the only type of morphological deformity; virtually all other morphological abnormalities are absent, iv) the presence of the deformity in the isolated gene pool within the radiating *Labeobarbus* assemblage, and v) the rapid growth of the abnormal individuals at 2-4 years of life compared to other sympatric morphs. Taking into account all the above, as well as the evidence for a genetic contribution to such abnormality in some other fish species, particularly the common carp, we assume that this phenomenon in this *Labeobarbus* population most likely has a genetic background. Moreover, the high-bodied phenotype in *Labeobarbus* may have a selective advantage as an anti-predator defense.

## Acknowledgments

Material for this study was collected within the scope of the Joint Ethiopian-Russian Biological Expedition (JERBE). We gratefully acknowledge the JERBE coordinators A.A. Darkov (Severtsov Institute of Ecology and Evolution, Moscow, Russia - IEE) and Simenew Keskes Melaku (Ministry of Innovation & Technology, Addis Ababa, Ethiopia) for administrative assistance. We express our gratitude to S.E. Cherenkov and Yu.Yu. Dgebuadze (both from IEE), Fekadu Tefera and Genanaw Tesfaye (both from National Fishery and Other Aquatic Life Research Center of the Ethiopian Institute of Agricultural Research - EIAR, Sebeta), M.V. Mina (Institute of Developmental Biology, Moscow, Russia) for sharing field operations and assistance in collecting material. We thank S.E. Cherenkov for his help with photography, F.N. Skil (IEE) for aging materials from experimentally reared barbs, O.N. Artaev for creating the map, M.V. Mina for critical comments on the manuscript, and Jacquelin DeFaveri for linguistic corrections. This work was financially supported by the Russian Science Foundation, project no. 19-14-00218.

## Supporting information

**S1 Table**. Occurrence of aberrant individuals or stocks with shortened vertebral column and abnormally short and high body in the different groups of fish. (DOCX)

**S2 Table**. Complete list of sampling sites. (XLSL)

**S3 Figure**. Photographs of two habitats at the main sampling (no. 1) of the middle Genale River – (A) the continuous pool and (B) the upper section of rapids downstream of the pool. (DOCX)

**S4 Table**. Size characteristics of the cast and gill net catches for different *Labeobarbus* forms from the middle Genale assemblage sampled in 2009 and 2019. Size characteristics of the aged individuals from the gill net catches of 2019. (DOCX)

**S5 Figure**. Variation in lower lip development in short individuals. (DOCX)

**S6 Figure**. Two piscivorous barbs with (A) normal and (B) deformed vertebral column. (DOCX)

**S7 Table**. Gill net catch composition and percentage of different *Labeobarbus* morphs in the middle Genale assemblage in 2009 and 2019. (DOCX)

**S8 Table**. Eigenvectors of the 10 most loaded characters from Fig. 4. (DOCX)

**S9 Table**. Data on vertebral counts. (DOCX)

**S10 Figure**. Supplement to Fig. 7. Distribution of deformed vertebrae along the vertebral column in individuals with 40 and 42 total vertebrae. (DOCX)

**S11 Figure**. Growth of short (SH), generalized (GN) and lipped (LP) forms in the middle Genale *Labeobarbus* assemblage sampled in 2009, estimated with standard length (SL). (DOCX)

